# The genetic architecture of osteoarthritis: insights from UK Biobank

**DOI:** 10.1101/174755

**Authors:** Eleni Zengini, Konstantinos Hatzikotoulas, Ioanna Tachmazidou, Julia Steinberg, Fernando P. Hartwig, Lorraine Southam, Sophie Hackinger, Cindy G. Boer, Unnur Styrkarsdottir, Daniel Suveges, Britt Killian, Arthur Gilly, Thorvaldur Ingvarsson, Helgi Jonsson, George C. Babis, Andrew McCaskie, Andre G. Uitterlinden, Joyce B. J. van Meurs, Unnur Thorsteinsdottir, Kari Stefansson, George Davey Smith, Mark J. Wilkinson, Eleftheria Zeggini

## Abstract

Osteoarthritis is a common complex disease with huge public health burden. Here we perform a genome-wide association study for osteoarthritis using data across 16.5 million variants from the UK Biobank resource. Following replication and meta-analysis in up to 30,727 cases and 297,191 controls, we report 9 new osteoarthritis loci, in all of which the most likely causal variant is non-coding. For three loci, we detect association with biologically-relevant radiographic endophenotypes, and in five signals we identify genes that are differentially expressed in degraded compared to intact articular cartilage from osteoarthritis patients. We establish causal effects for higher body mass index, but not for triglyceride levels or type 2 diabetes liability, on osteoarthritis.

---

Osteoarthritis (OA) is the most prevalent musculoskeletal disease and the most common form of arthritis^1^. The hallmarks of OA are degeneration of articular cartilage, remodelling of the underlying bone and synovitis^2^. A leading cause of disability worldwide, it affects 40% of individuals over the age of 70 and is associated with an increased risk of comorbidity and death^3,4^. The health economic burden of OA is rising, commensurate with longevity and obesity rates, and there is currently no curative therapy^5,6^. The heritability of OA is ∼50%^7-9^, and previous genetic studies have identified 21 loci in total^10^, traversing hip, knee and hand OA with limited overlap^9,11-25^. Here we conduct the largest OA genome wide association study (GWAS) to date, using genotype data across 16.5 million variants from UK Biobank. We define OA based on both self-reported status and through linkage to Hospital Episode Statistics data, and on joint-specificity of disease (knee and/or hip) (Supplementary Fig. 1).

## RESULTS

### Disease definition and power to detect genetic associations

We compare and contrast the hospital diagnosed (n=10,083 cases) to self-reported (n=12,658 cases) OA GWAS drawn from the same UK Biobank dataset (with non-OA controls selected to be ∼4x the number of cases to preserve power for common alleles while avoiding case:control imbalance causing association tests to misbehave for low frequency variants^26^) (Supplementary Tables 1-3, Supplementary Figs. 2-4; Methods). We find power advantages with the self-reported dataset, indicating that the increase in sample size overcomes the limitations associated with phenotype uncertainty. When evaluating the accuracy of disease definition, we find that self-reported OA has modest positive predictive value (PPV=30%) and sensitivity (37%), but a high negative predictive value of 95% and high specificity, correctly identifying 93% of individuals who do not have OA (Supplementary Table 4). In terms of power to detect genetic associations, the self-reported OA dataset has clear advantages commensurate with its larger samples size (Figure 1). For example, for a representative complex disease-associated variant with minor allele frequency (MAF) 30% and allelic odds ratio 1.10, the self-reported and hospital diagnosed OA analyses have 80% and 56% power to detect an effect at genome-wide significance (i.e., *P* <5.0×10^−8^), respectively (Supplementary Table 5).

**Figure 1.**
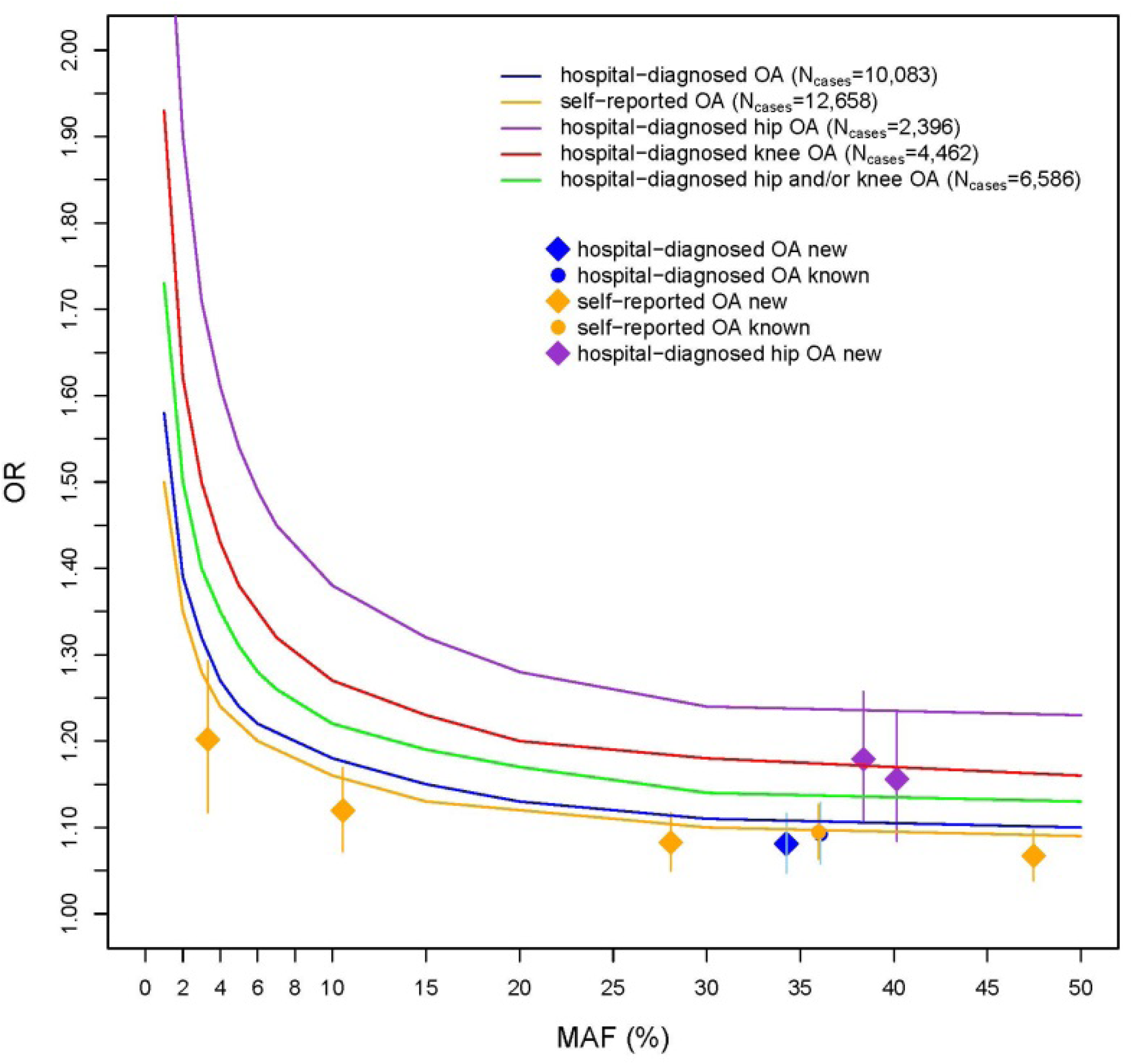
Power to detect association in the discovery stage. Odds ratios (ORs) and 95% confidence intervals as a function of minor allele frequencies (MAF). Newly reported variants are denoted in diamonds, while known variants are denoted in circles. The curves indicate 80% power at the genome-wide significance threshold of *P*<5.0×10^−8^, for the number of cases and controls of each trait at the discovery stage.

We find nominally significant evidence for concordance between the direction of effect at previously reported OA loci and the discovery analyses for hospital diagnosed OA definitions (Supplementary Table 6 and Supplementary note), indicating that a narrower definition of disease may provide better effect size estimates albeit limited by power to identify robust statistical evidence for association.

The only previously established OA signal to reach genome-wide significance in the discovery dataset was rs143383 in *GDF5* (Supplementary Table 7 and Supplementary note), identified as genome-wide significant in the self-reported OA analysis (OR(95%CI) 0.91(0.89-0.94), *P*=5.11×10^−9^), but not in the hospital diagnosed OA analysis (0.91(0.89-0.94), *P*=3.53×10^−7^), although the effect sizes are clearly very similar. rs143383 remained genome-wide significantly associated with self-reported OA when a random subset of the dataset was taken to match the sample size of the hospital diagnosed OA dataset (sensitivity analysis: 0.90(0.88-0.94), *P*=1.55×10^−8^; Methods).

### Heritability estimates and genetic correlation between OA definitions

We find that common-frequency variants explain 12% of OA heritability when using self-reported status, and 16% when using hospital records (19% of hip OA and 15% of knee OA heritability) (Supplementary Table 8). Heritability estimates from self-reported and hospital records were not different from each other (*P*=0.06 and *P*=0.07 when using the Pearson’s product-moment correlation or the Spearman’s rank correlation, respectively) (Supplementary Table 9). The concordance between self-reported and hospital diagnosed OA was further substantiated by the high genetic correlation estimate of the two disease definitions (87%, *P*=3.14×10^−53^) (Supplementary Table 10). We find strong genome-wide correlation between hip OA and knee OA (88%, *P*=1.96×10^−6^), even though previously reported OA loci are predominantly not shared between the two OA joint sites. Based on this new observation of a substantial shared genetic aetiology, we sought replication of association signals across joint sites.

### Identification of novel OA loci

We took 173 variants with *P*<1.0×10^−5^ and MAF>0.01 forward to replication in an Icelandic cohort of up to 18,069 cases and 246,293 controls (Supplementary Fig. 1, Supplementary Tables 11-15; Methods). Following meta-analysis in up to 30,727 cases and 297,191 controls, we report seven genome-wide significant associations at novel loci, and two further new replicating signals just below the genome-wide significance threshold (*P*<6.0×10^−8^; Table 1, Figure 2).

**Figure 2.**
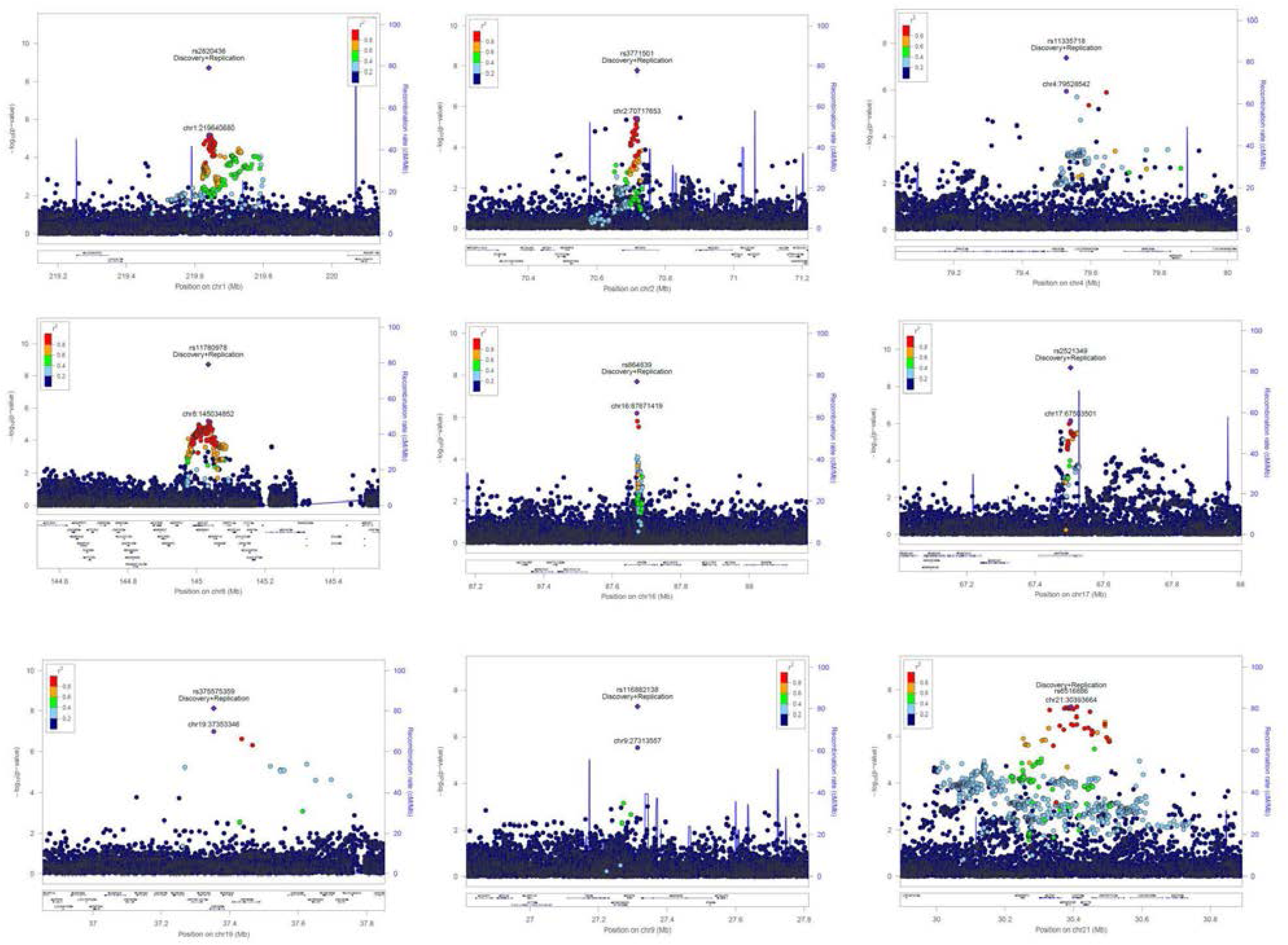
Regional association plots for the nine novel OA loci. The y axis represents the negative logarithm (base 10) of the variant P-value and the x axis represents the position on the chromosome, with the name and location of genes and nearest genes shown in the bottom panel. The variant with the lowest P-value in the region after combined discovery and replication is marked by a purple diamond. The same variant is marked by a purple dot showing the discovery P-value. The colours of the other variants indicate their r2 with the lead variant.

**Table 1.** Association summary statistics for the nine novel replicating signals. ^a^The number of cases required for 80% power is calculated on the basis of the replication study OR estimate, sample size-weighted effect allele frequency across the discovery and replication studies, and assuming 4 controls per case. EA: effect allele; EAF: effect allele frequency; OR: odds ratio.

| rsID | nearest gene(s) | EA | discovery phenotype | discovery EAF | discovery OR (95% CI) | discovery *P*-value | discovery n cases/controls | replication phenotype | replication EAF | replication OR (95% CI) | replication *P*-value | replication n cases/controls | overall OR (95% CI) | overall *P*-value | overall n cases/controls | number of cases for 80% power at $\alpha=0.05^a$ | number of cases for 80% power at $\alpha=5 \times 10^{-8}^a$ |
| --- | --- | --- | --- | --- | --- | --- | --- | --- | --- | --- | --- | --- | --- | --- | --- | --- | --- |
| rs2820436 | *SLC30A10*,<br>*LYPLAL1* | C | Hospital diagnosed OA | 0.66 | 0.92<br>(0.9-0.96) | $6.45 \times 10^{-6}$ | 10,083/<br>40,425 | OA at any site | 0.64 | 0.94<br>(0.91-0.97) | $8.71 \times 10^{-5}$ | 18,069/<br>246,293 | 0.93<br>(0.91-0.96) | $2.01 \times 10^{-9}$ | 28,152/<br>286,664 | 5,572 | 28,115 |
| rs3771501 | *TGFA* | G | Self-reported OA | 0.53 | 0.94<br>(0.91-0.96) | $3.81 \times 10^{-6}$ | 12,658/<br>50,898 | OA at any site | 0.54 | 0.95<br>(0.92-0.98) | 0.001069 | 18,069/<br>246,293 | 0.94<br>(0.92-0.96) | $1.66 \times 10^{-8}$ | 30,727/<br>297,137 | 7,474 | 37,709 |
| rs11335718 | *ANXA3* | A | Self-reported OA | 0.11 | 1.12<br>(1.07-1.17) | $1.12 \times 10^{-6}$ | 12,658/<br>50,898 | Knee OA | 0.11 | 1.1<br>(1.02-1.2) | 0.014675 | 4,672/<br>172,791 | 1.11<br>(1.07-1.16) | $4.26 \times 10^{-8}$ | 17,330/<br>223,689 | 5,278 | 26,629 |
| rs11780978 | *PLEC* | A | Hospital diagnosed hip | 0.4 | 1.16<br>(1.08-1.23) | $6.24 \times 10^{-6}$ | 2,396/<br>9,593 | Hip OA | 0.39 | 1.11<br>(1.051-1.16) | $4.55 \times 10^{-5}$ | 5,714/<br>199,421 | 1.13<br>(1.08-1.17) | $1.98 \times 10^{-9}$ | 8,083/<br>209,014 | 1,870 | 9,435 |
| rs116882138 | *MOB3B*,<br>*EQTN* | A | Hospital diagnosed hip and/or knee OA | 0.02 | 1.4<br>(1.22-1.6) | $2.96 \times 10^{-6}$ | 6,586/<br>26,384 | Knee OA | 0.02 | 1.27<br>(1.07-1.5) | 0.006552 | 4,672/<br>172,791 | 1.34<br>(1.21-1.49) | $5.09 \times 10^{-8}$ | 11,258/<br>199,175 | 6,056 | 30,556 |
| rs2521349 | *MAP2K6* | A | Hospital diagnosed hip OA | 0.38 | 1.18<br>(1.11-1.26) | $6.85 \times 10^{-7}$ | 2,396/<br>9,593 | Hip OA | 0.37 | 1.1<br>(1.05-1.16) | 0.000103 | 5,714/<br>199,421 | 1.13<br>(1.09-1.18) | $9.95 \times 10^{-10}$ | 8,083/<br>209,014 | 3,818 | 19,265 |
| rs864839 | *JPH3* | T | Self-reported OA | 0.72 | 1.08<br>(1.05-1.12) | $6.21 \times 10^{-7}$ | 12,658/<br>50,898 | Hip OA | 0.7 | 1.07<br>(1.02-1.13) | 0.008275 | 5,714/<br>199,421 | 1.08<br>(1.05-1.11) | $2.01 \times 10^{-8}$ | 18,372/<br>250,319 | 5,297 | 26,724 |
| rs375575359 | *ZNF345* | C | Self-reported OA | 0.03 | 1.2<br>(1.12-1.29) | $9.96 \times 10^{-8}$ | 12,658/<br>50,898 | Knee OA | 0.05 | 1.15<br>(1.02-1.29) | 0.025177 | 4,672/<br>172,791 | 1.21<br>(1.14-1.3) | $7.54 \times 10^{-9}$ | 17,330/<br>223,689 | 2,285 | 11,529 |
| rs6516886 | *RWDD2B*,<br>*USP16*,<br>*LTN1* | T | Hospital diagnosed hip and/or knee OA | 0.75 | 1.13<br>(1.08-1.19) | $5.36 \times 10^{-8}$ | 6,586/<br>26,384 | Hip OA | 0.76 | 1.06<br>(1-1.12) | 0.055135 | 5,714/<br>199,421 | 1.1<br>(1.06-1.14) | $5.84 \times 10^{-8}$ | 12,300/<br>225,805 | 7,844 | 39,577 |

We identify association between rs2521349 and hip OA (OR 1.13 (95% CI 1.09-1.17), *P*= 9.95×10^−10^, effect allele frequency [EAF] 0.37). rs2521349 resides in an intron of *MAP2K6* on chromosome 17. *MAP2K6* is an essential component of the p38 MAP kinase mediated signal transduction pathway, involved in various cellular processes in bone, muscle, fat tissue homeostasis and differentiation^27^. The MAPK signalling pathway has been closely associated with osteoblast differentiation, chondrocyte apoptosis and necrosis, and reported to be differentially expressed in OA synovial tissue samples^28-38^. In animal model studies, its activity has been found to be important in maintaining cartilage health and it has been proposed as a potential OA diagnosis and treatment target^33,39,40^. rs11780978 on chromosome 8 is also associated with hip OA with a similar effect size (OR 1.13 (95% CI 1.08-1.17), p=1.98×10^−9^, EAF 0.39). This variant is located in the intronic region of the plectin gene, *PLEC*. We find rs11780978 to be nominally associated with the radiographically derived endophenotype of minimal joint space width (beta -0.0291, SE 0.0129, *P*=0.024) (Table 2; Methods). The direction of effect is consistent with the established clinical association between joint space narrowing and OA. *PLEC* encodes plectin, which acts as a link between components of the cytoskeleton as a dynamic organizer of intermediate filament networks^41^. Plectin has been found to be downregulated in OA-affected meniscus compared with healthier tissue and is reported to play a key role in causing dystrophic changes in muscle^42,43^. Functional studies in mice have shown an effect on skeletal muscle tissue and correlation with decreased body weight, size and postnatal growth^44^. The chromosomal region surrounding rs11780978 also contains correlated variants (r^2^ 0.78-1) associated with metabolic and anthropometric traits in humans.

**Table 2.**
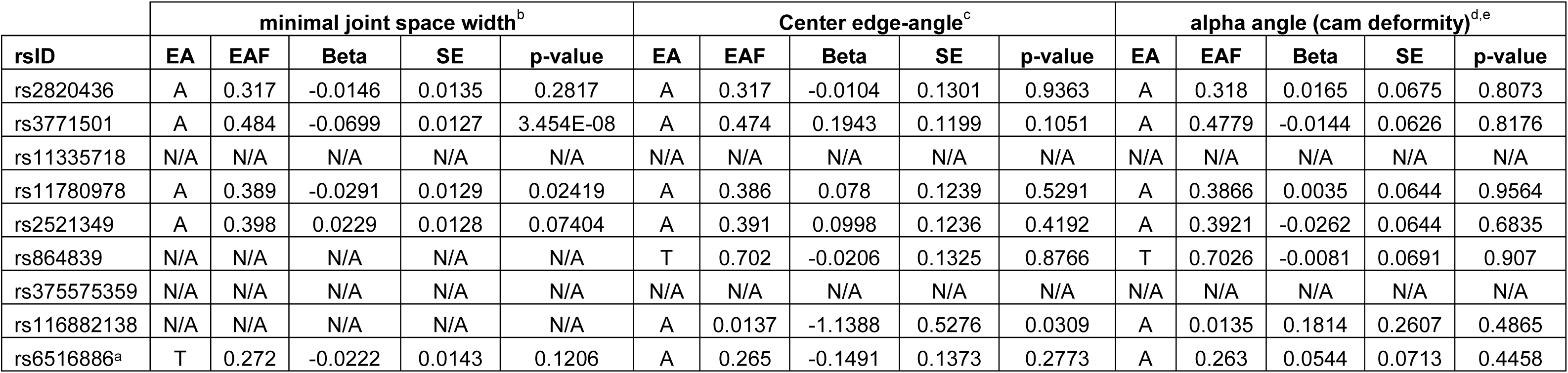
Association of the 9 novel OA loci with radiographically-derived OA endophenotypes. ^a^For minimal joint space width, proxy variant rs2150403 (r2=0.99 with rs6516886) was used. ^b^Sample size=13,013. ^c^Sample size=6,880. ^d^Sample size cases=639. ^e^Sample size controls=4,339. EA: effect allele; EAF: effect allele frequency; SE: standard error; N/A: not available.

rs2820436 is an intergenic variant located 24kb upstream of long non-coding RNA *RP11-392O17.1* and 142kb downstream of zinc finger CCCH-type containing 11B pseudogene *ZC3H11B*, and is associated with OA across any joint site (OR 0.93 (95% CI 0.91-0.95), *P*=2.01x10^−9^, EAF 0.65). It also resides within a region with multiple metabolic and anthropometric trait-associated variants^45,46^, with which it is correlated (r^2^ 0.18-0.88). rs375575359 resides in an intron of the zinc finger protein 345 gene, *ZNF345,* on chromosome 19. It was prioritised based on OA at any joint site and is more strongly associated with knee OA in the replication dataset (OR 1.21 (95% CI 1.14-1.30), *P*=7.54×10^−9^, EAF 0.04). Similarly, rs11335718 on chromosome 4 was associated with OA in the discovery and knee OA in the replication stage (OR 1.11 (95% CI 1.07-1.16), p= 4.26×10^−8^, EAF 0.10). rs11335718 is an intronic variant in the annexin A3 gene, *ANXA3*. By meta-analysing the any site OA phenotype across the discovery and replication datasets, we report *P*=2.6×10^−5^ and *P*=1.32×10^−7^ for rs375575359 and rs11335718, respectively (Supplementary Table 11). A recent mouse model study supports the involvement of a similar motif zinc finger protein expression (ZFP36L1) with osteoblastic differentiation^47^. rs3771501 (OR 0.94 (95% CI 0.92-0.96), *P*=1.66×10^−8^, EAF 0.53) is associated with OA at any site and resides in an intron of the transforming growth factor alpha gene, *TGFA*. *TGFA* encodes an epidermal growth factor receptor ligand and is an important integrator of cellular signalling and function^48,49^. We detect association of rs3771501 with minimal joint space width (beta -0.0699, SE 0.0127, *P*=3.45×10^−8^) (Table 2; Methods), i.e. the OA risk increasing allele is also associated with decreased joint cartilage thickness in humans. Variation in this gene has previously been associated with cartilage thickness, suggestively associated with hip OA, and found to be differentially expressed in OA cartilage lesions compared to non-lesioned cartilage^50^. Functional studies have revealed that TGFA regulates the conversion of cartilage to bone during the process of endochondral bone growth, that it is a dysregulated cytokine present in degrading cartilage in OA and a strong stimulator of cartilage degradation upregulated by articular chondrocytes in experimentally induced and human OA^51-54^. The function of *TGFA* has also been associated with craniofacial development, palate closure and decreased body size^48^. rs864839 resides in the intronic region of the junctophilin 3 gene, *JPH3*, on chromosome 16, and was discovered based on the any joint site OA analysis. It is more strongly associated with hip OA in the replication dataset (OR 1.08 (95% CI 1.05-1.11), *P*=2.1×10^−8^, EAF 0.71). By meta-analysing the any site OA phenotype across the discovery and replication datasets, we report *P*=7.02×10^−6^ (Supplementary Table 11). JPH3 is involved in the formation of junctional membrane structure, regulates neuronal calcium flux and is reported to be expressed in pancreatic beta cells and in the regulation of insulin secretion^55,56^.

We detect two further replicating signals that narrowly fail to reach the genome-wide significance threshold. rs116882138 is most strongly associated with hip and/or knee OA in the discovery and with knee OA in the replication dataset (OR 1.34 (95% CI 1.21-1.49), *P*=5.09×10^−8^, EAF 0.02). It is an intergenic variant located 11kb downstream of the kinase activator 3B gene, *MOB3B*, and 16kb upstream of the equatorin, sperm acrosome associated gene, *EQTN*, on chromosome 9. We find rs116882138 to be nominally associated with acetabular dysplasia as measured with Center Edge-angle (beta -1.1388, SE 0.5276, *P*=0.031) (Table 2; Methods).

Finally, rs6516886 was prioritised based on the hip and/or knee OA discovery analysis and is more strongly associated in the hip OA replication dataset (OR 1.10 (95% CI 1.06-1.14), p=5.84×10^−8^, EAF 0.75). rs6516886 is situated 1kb upstream of the RWD domain containing 2B gene, *RWDD2B*, on chromosome 21. *LTN1* (listerin E3 ubiquitin protein ligase 1), a protein coding gene which resides at a distance of 28kb from the variant, has been reported to affect musculoskeletal development in a mouse model^57^.

### Functional analysis

We tested whether coding genes within 1Mb surrounding the novel OA-associated variants were differentially expressed at 1% false discovery rate (FDR) in chondrocytes extracted from intact compared to degraded cartilage from OA patients undergoing total joint replacement surgery using molecular phenotyping through quantitative proteomics and RNA sequencing (Table 3; Methods).

**Table 3.** Genes in the novel OA-associated signals with significantly different gene expression and/or protein abundance in intact compared to degraded articular cartilage. logFC: log2-fold change (increase means higher value in degraded cartilage); FDR: false discovery rate; OA: Osteoarthritis.

| Index variant | Gene | Position<br>(chr:start-end) | Distance from<br>index variant (kb) | Protein<br>logFC | Protein<br>FDR *q-value* | RNAseq<br>logFC | RNAseq<br>FDR *q-value* |
| --- | --- | --- | --- | --- | --- | --- | --- |
| rs3771501 | *PCYOX1* | 2:70484518- 70508323 | 209.3 | -0.27 | 0.0042 | 0.27 | 0.0047 |
| rs3771501 | *FAM136A* | 2:70523107- 70529222 | 188.4 | - | - | -0.20 | 0.0066 |
| rs6516886 | *BACH1* | 21:30566392- 31003071 | 172.7 | - | - | 0.32 | 0.0019 |
| rs6516886 | *MAP3K7CL* | 21:30449792-30548210 | 56.1 | - | - | 0.41 | 0.0021 |
| rs11780978 | *BOP1* | 8:145486055-145515082 | 451.2 | - | - | -0.26 | 0.0030 |
| rs116882138 | *PLAA* | 9:26904081-26947461 | 366 | - | - | 0.20 | 0.0027 |
| rs375575359 | *ZNF382* | 19:37095719-37119499 | 233.8 | - | - | 0.39 | 0.0031 |

*PCYOX1*, located 209kb downstream of rs3771501, showed significant evidence of differential expression (1.21-fold higher post-normalisation in degraded cartilage at the RNA level, *q-value*=0.0047; and 1.17-fold lower abundance at the protein level, *q*=0.0042). The protein product of this gene catalyses the degradation of prenylated proteins^58^. *PCYOX1* has been reported to be overexpressed in human dental pulp derived osteoblastoids compared to osteosarcoma cells^59^. Further investigation into the chondrocyte and peripheral secretome is warranted to assess the potential of this molecule as a biomarker for OA progression. *FAM136A*, located 188kb upstream of the same variant (rs3771501), showed 1.13-fold lower transcriptional levels in chondrocytes from degraded articular cartilage (*q*=0.0066).

*BACH1* and *MAP3K7CL*, located in the vicinity of rs6516886, showed evidence of differential expression (1.26-fold, *q*=0.0019, and 1.37-fold, *q*=0.0021, higher transcription in degraded tissue, respectively). BACH1 is a transcriptional repressor of Heme oxygenase-1 (HO-1). Studies in Bach1 deficient mice have independently suggested inactivation of Bach1 as a novel target for the prevention and treatment of meniscal degeneration^60^ and of osteoarthritis^61^.

Finally, *PLAA* and *ZNF382* located proximal to rs116882138 and rs375575359, respectively, showed higher transcription levels in degraded compared to intact cartilage (1.15-fold, *q*=0.0027, and 1.31-fold, *q*=0.0031, respectively). *BOP1* located 451kb downstream of rs11780978 showed 1.17-fold lower levels of transcription in degraded tissue (*q*=0.003).

### Fine-mapping points to non-coding variants at all novel loci

For five of the new loci, the sum of probabilities of causality of all variants in the fine-mapped region was ≥0.95 (>0.99 for two signals), and for two further loci it was >0.93 (Supplementary Table 16; Methods). The majority of variants within each credible set have marginal posterior probabilities, while only a small number of variants have posterior probability of association (PPA) >0.1; these account for 25-92% of PPA across the different regions. The credible set of four signals can be narrowed down to 3 variants, one signal to 2 variants, and one signal to 1 variant with a probability of causality >0.1. For all 9 regions the variant identified as the most likely to be causal is non-coding (Supplementary Table 17).

Although the correlation between the different functional annotation scores used is variable (Supplementary Fig. 5), the Pearson’s correlation coefficient of the posterior probability of association of variants based on different annotations in the fine-mapped regions is 1, and therefore we find that the choice of *in silico* functional prediction scores had limited impact on the fine-mapping results.

For three of the regions, the credible set variants with PPA>0.1 were contained within a single gene, although we cannot draw inferences that these genes be causal. The signal indexed by rs2521349 was fine-mapped to 33 variants spanning two genes, including two variants with PPA>0.1 within 153 base pairs of each other in *MAP2K6*. This narrow chromosomal region is over 200 times smaller than the original association peak (as delineated by variants with r^2^>0.5 with the index variant). Similarly, the fine-mapped region surrounding rs3771501 contains 33 variants traversing 4 genes across 147kb. The credible set contains three variants with probability of causality >0.1, all within 6.2kb in the second intron of *TGFA*. Finally, rs864839 was identified as the most likely causal variant with a high probability of 0.55. The 95% credible interval for this region contains 250 variants across 58kb, all within *JPH3*, including three variants with a probability of causality >0.1 and a combined PPA of >91%, spanning a narrower interval of 5.2kb within the first intron of the gene. One signal was fine-mapped to a single variant, rs116882138, with 70% PPA, with the credible interval reaching 75% probability of at least one causal variant.

### Gene-based analyses

Gene set analysis identified *UQCC1* and *GDF5*, located in close vicinity to each other on chromosome 20, as key genes with consistent evidence for significant association with OA across phenotype definitions (Supplementary Table 18 and Supplementary note). *GDF5* codes for growth differentiation factor 5, a member of the TGFbeta superfamily, and plays a central role in skeletal development. Pathway analyses identified significant associations between self-reported OA and anatomical structure morphogenesis (*P*=4.76×10^−5^), ion channel transport (*P*=8.98×10^−5^); hospital diagnosed hip OA and activation of MAPK activity (*P*=1.61×10^−5^); hospital diagnosed knee OA and histidine metabolism (*P*=1.02×10^−5^); and between hospital diagnosed hip and/or knee OA and recruitment of mitotic centrosome proteins and complexes (*P*=8.88×10^−5^) (Supplementary Table 19, Supplementary Fig. 6).

### Genetic links between OA and epidemiologically linked traits

Established clinical risk factors for OA include increasing age, female sex, obesity, occupational exposure to high levels of joint loading activity, previous injury, smoking status and family history of OA. We estimated the genome-wide genetic correlation between OA and 219 other traits and diseases and identified 35 phenotypes with significant (5% FDR) genetic correlation with OA across definitions, with large overlap between the identified phenotypes (Supplementary Fig. 7, Figure 3, Supplementary Table 20; Methods).

**Figure 3.**
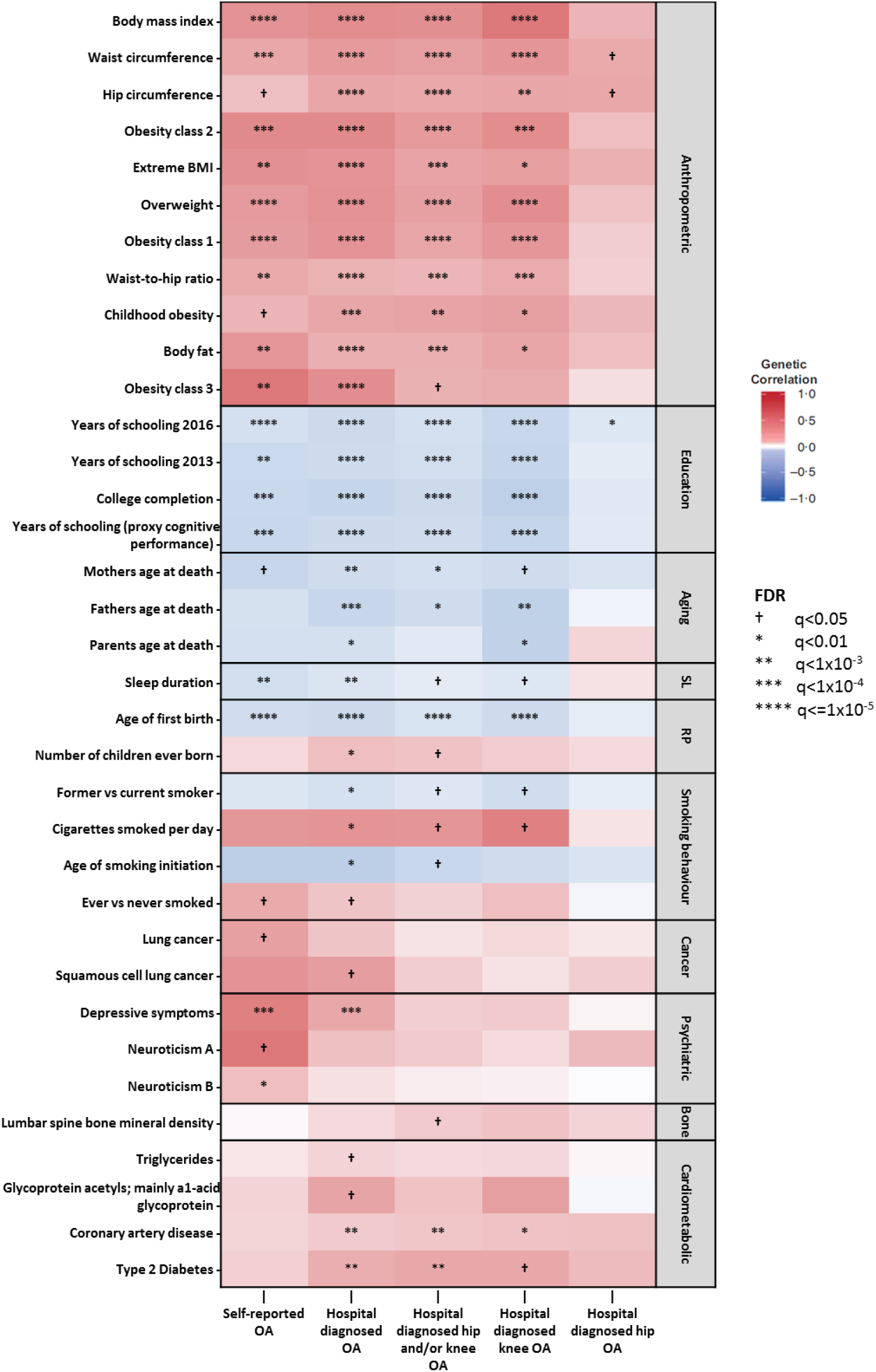
Heat map of genetic correlations between OA phenotypes in UK Biobank and 35 traits grouped in 10 categories from GWAS consortia. Symbols and hues depict the FDR *q-values* and strength of the genetic correlation (darker shade denotes stronger correlation), respectively. Red and blue indicate positive and negative correlations, respectively. RP: reproductive; SL: sleep.

The phenotypes with significant genetic correlations (rg) fall into the following broad categories: obesity, BMI and related anthropometric traits (rg>0); type 2 diabetes (rg>0); educational achievement (rg<0); neuroticism, depressive symptoms (rg>0), and sleep duration (rg<0); mother’s, father’s, or parents’ age at death (rg<0); reproductive phenotypes, including age at first birth (rg<0) and number of children ever born (rg>0); smoking, including age of smoking initiation (rg<0) and ever smoker (rg>0), and lung cancer (rg>0) (Figure 3, Supplementary Table 20). The four phenotypes with significant genetic correlation in all analyses were: years of schooling, waist circumference, hip circumference and BMI.

We find a nominally significant positive genetic correlation with rheumatoid arthritis, which did not pass multiple-testing correction for self-reported and hospital diagnosed OA (rg=0.14-0.19, FDR 10%-12%). Of musculoskeletal phenotypes, lumbar spine bone mineral density showed positive genetic correlation with hospital diagnosed hip and/or knee OA (rg=0.2, FDR=3%) but did not reach significance in other analyses.

### Disentangling causality

We undertook Mendelian randomisation (MR) analyses^62^ to strengthen causal inference regarding modifiable exposures that could influence OA risk (Supplementary Tables 21-24; Methods). Each kg/m^2^ increment in body mass index was predicted to increase risk of self-reported OA by 1.11 (95% CI: 1.07-1.15, *P*= 8.3x10^−7^). This result was consistent across MR estimators (OR ranging from 1.52 to 1.66) and disease definition (OR ranging from 1.66 to 2.01). Consistent results were also observed for other obesity-related measures, such as waist (OR: 1.03 per cm increment; 95% CI: 1.02-1.05, *P*=5×10^−4^) and hip circumference (OR: 1.03 per cm increment; 95% CI: 1.01-1.06, *P*=0.021). OR for type 2 diabetes liability and triglycerides were consistently small in magnitude across estimators and OA definitions; given that analyses involving those traits were well-powered (Supplementary Table 25), these results are compatible with either a weak or no causal effect. Results for years of schooling were not consistent across estimators, and there was evidence of directional horizontal pleiotropy, thus hampering any causal interpretation (Figure 4). There was some evidence for a site-specific causal effect of height on knee OA (OR: 1.13 per standard deviation increment; 95% CI: 1.02-1.25, *P*=0.023), which was consistent across estimators. One-sample MR analyses corroborated these findings, with only obesity-related phenotypes presenting strong statistical evidence after multiple testing correction (Supplementary Table 26). These analyses did not detect reliable effects of smoking or reproductive traits on osteoarthritis (Supplementary Tables 27 and 28).

**Figure 4.**
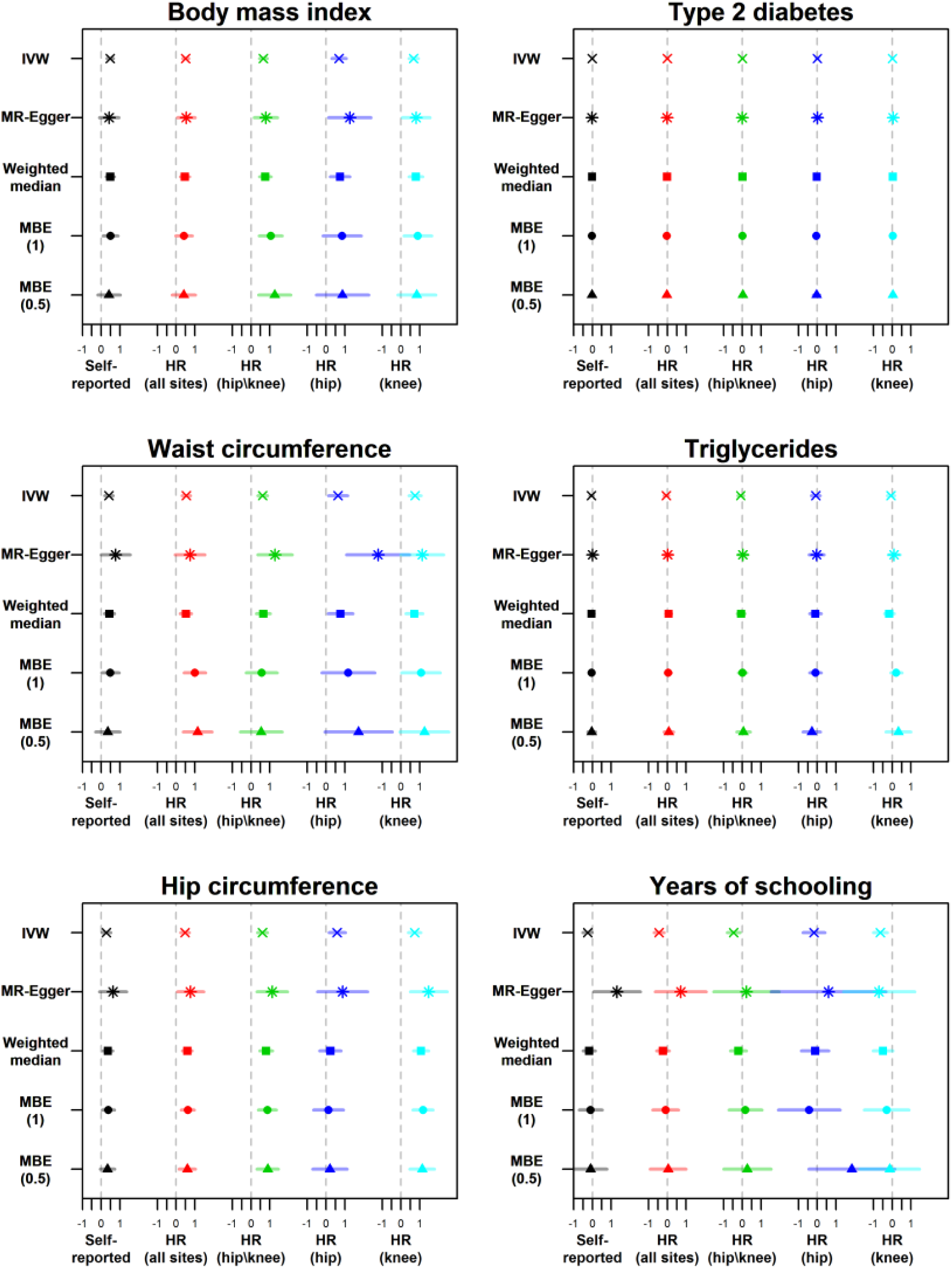
Two-sample Mendelian randomization estimates of the effect of obesity-related measures, triglyceride levels, years of schooling (in standard deviations units) and type 2 diabetes liability (in ln(odds ratio) units) on different definitions of osteoarthritis. HR: hospital record. IVW: inverse-variance weighting. MBE: mode-based estimate. MBE (1): tuning parameter *ψ*=1. MBE (0.5): tuning parameter *ψ*=0.5.

## DISCUSSION

In order to improve our understanding of the genetic aetiology of osteoarthritis, we have conducted a study combining genotype data in up to 327,918 individuals. We identify 9 novel, robustly replicating loci associated with OA, two of which fall just under the genome-wide significance threshold; this constitutes a substantial increase in the number of known OA loci. Taken together, all established OA loci account for 26.3% of trait variance (Supplementary Fig. 8). The key attributes of this study are sample size and the homogeneity of the UK Biobank dataset, coupled with independent replication, independent association with clinically-relevant radiographic endophenotypes, and functional genomics follow up in primary osteoarthritis tissue. We have further capitalised on the wealth of available genome-wide summary statistics across complex traits to identify genetic correlations between OA and multiple molecular, physiological and behavioural phenotypes, followed by formal Mendelian randomization analyses to assess causality and disentangle complex cross-trait epidemiological relationships.

The vast majority of novel signals are at common frequency variants and confer small to modest effects, in line with a highly polygenic model underpinning OA risk. We identify one low-frequency variant association with OA (MAF 0.02) with a modest effect size (combined OR 1.34). Even though well-powered to detect them, we find no evidence for a role of low frequency variation of large effect in OA susceptibility (Supplementary Table 5). We estimate the requirement of up to 40,000 OA cases and 160,000 controls to recapitulate the effects identified in this study at genome-wide significance based on the sample size-weighted effect allele frequencies and replication cohort odds ratio estimates (Table 1).

We integrated functional information with statistical evidence for association to fine-map the location of likely causal variants and genes. All predicted most likely causal variants reside in non-coding sequence: 6 are intronic and 3 are intergenic. We are able to refine the association signal to a single variant in one occasion, and to variants residing within a single gene in three instances, although the mechanism of action could be mediated through other genes in the vicinity.

We empirically find self-reported OA definition to be a powerful tool for genetic association studies, for example as evidenced by the fact that the established *GDF5* OA locus reaches genome-wide significance in the self-reported disease status analyses only. Furthermore, our formal evaluation of phenotype accuracy showing low sensitivity and high specificity is consistent with a recent report comparing self-reported data with hospitalisation records for heart disease, stroke, bronchitis, depression and cancer in the UK Biobank resource^63^. Hospital diagnosed data for OA are potentially incomplete; as patients with OA are usually hospitalized by the time OA is severe and has to be treated by joint replacement surgery. Several published epidemiological studies have studied OA via self-report^64-69^ and validation of self-reported status against primary care records has yielded similar conclusions^66^. We also find very high genetic correlation between self-reported and hospital diagnosed OA, and similar SNP heritability estimates, corroborating the validity of self-reported OA status for genetic studies.

We identify strong genome-wide correlation between hip and knee OA, indicating a substantial shared genetic aetiology that has been hitherto missed. We therefore sought replication of signals across joint sites and identify multiple instances of cross-joint site validation. In all of these instances, signals were detected in the larger discovery analysis of OA at any site, and independently replicated in joint-specific definitions of disease. Further analysis in larger sample sets with precise phenotyping will help distinguish signal specificity.

Two of the newly identified signals, indexed by rs11780978 and rs2820436, reside in regions with established metabolic and anthropometric trait associations^45,70,71^. OA is epidemiologically associated with increased BMI, and the association is stronger for knee OA. In line with this, we find higher genetic correlation between BMI and knee OA (rg=0.52, *P*=2.2×10^−11^), than with hip OA (rg=0.28, *P*=4×10^−4^). BMI is also known to be genetically correlated with education phenotypes, depressive symptoms, reproductive and other phenotypes; hence, some of the genetic correlations for OA observed here could be mediated through BMI. However, for the education and personality/psychiatric phenotypes, the strength of genetic correlations observed here for OA is substantially higher than the genetic correlations observed for BMI (e.g. hospital diagnosed OA and years of schooling rg=-0.45, *P*=5×10^−27^, while BMI and years of schooling have rg=-0.27, *P*=9×10^−32^; hospital diagnosed OA and depressive symptoms rg=0.49, *P*=6×10^−7^, while BMI and depressive symptoms have rg=0.10, *P*=0.023). Epidemiologically, lower educational levels are known to be associated particularly with risk of knee OA, even when adjusting for BMI^72^.

Mendelian randomization provided further insight into the nature of the genetic correlations we observed. In the case of BMI and other obesity-related measures, there was evidence for a causal effect of those phenotypes on osteoarthritis. This result corroborates the findings from conventional observational studies^73-75^, which are prone to important limitations (such as reverse causation and residual confounding) regarding causal inference^76,77^. For all other exposure phenotypes, there was no convincing evidence for a causal effect on OA risk, suggesting that the genetic correlations detected by LD score regression may be mostly due to horizontal pleiotropy, although for some phenotypes the MR analyses were underpowered (Supplementary Table 25). In the case of triglycerides and liability to type 2 diabetes, the Mendelian randomization analyses had sufficient power to rule out non-small causal effects, suggesting that these phenotypes have at most weak effects on OA risk. These findings suggest that associations typically seen in conventional observational epidemiological studies^78-82^ are likely due to biases that were not measured or appropriately accounted for. In the case of educational attainment, lack of consistency across estimators hampered any causal inference from MR, indicating that other strategies to interrogate causality are required^83^.

Crucially, structural changes in the joint usually precede the onset of symptoms for OA. Articular cartilage is an avascular, aneural tissue. It provides tensile strength, compressive resilience and a low-friction articulating surface. Chondrocytes are the only cell type in cartilage. The accessibility of relevant tissue at joint replacement surgery enables the deployment of multi-omics to dissect the molecular disease processes in the relevant cells. The mode of function of non-coding DNA is linked to context-dependent gene expression regulation, and identification of the causal variants and the genes they affect requires experimental analysis of genome regulation in the right cell type. Our functional analysis of genes in OA-associated regions and pathways identified differentially expressed molecules in chondrocytes extracted from degraded compared to intact articular cartilage. Cartilage degeneration is a key hallmark of OA pathogenesis and regulation of these genes could be implicated in disease development and progression.

Osteoarthritis is a leading cause of disability worldwide and carries a substantial public health and health economics burden. Here, we have gleaned novel insights into the genetic aetiology of OA, and have implicated genes with translational potential^33,39,40,60,61^. Going forward, large-scale whole genome sequencing studies of well-phenotyped individuals across diverse populations will capture the full allele frequency and variation type spectrum, and afford us further insights into the causes of this debilitating disease.

## ACKNOWLEDGEMENTS

This research has been conducted using the UK Biobank Resource under Application Number 9979. This work was funded by the Wellcome Trust (WT098051). We are grateful to Roger Brooks, Jyoti Choudhary and Theodoros Roumeliotis for their contribution to the functional genomics data collection.

## DATA AVAILABILITY

All RNA sequencing data have been deposited to the European Genoome/Phenome Archive (cohort 1: EGAD00001001331; cohort 2: EGAD00001003355; cohort 3: EGAD00001003354).

## ONLINE METHODS

### Accuracy of self-reported data

We evaluated the classification accuracy of self-reported disease status by estimating the sensitivity, specificity, positive predictive values (PPV) and negative predictive values (NPV) in the self-reported and hospital diagnosed disease definition datasets. We performed a sensitivity test to evaluate the true positive rate by calculating the proportion of individuals diagnosed with OA that were correctly identified as such in the self-reported analysis, and a specificity test to evaluate the true negative rate by calculating the proportion of individuals not diagnosed with OA that were correctly identified as such in the control set. The number of individuals overlapping between the self-reported (n_SR_=12,658) and hospital-diagnosed (n_HD_=10,083) datasets was n_OVER_=3,748. The total number of individuals was n_TOT_=138,993.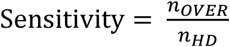; 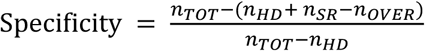; 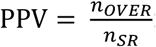; 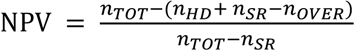.

### Discovery GWAS

UK Biobank’s scientific protocol and operational procedures were reviewed and approved by the North West Research Ethics Committee (REC Reference Number: 06/MRE08/65). The 1st UK Biobank release of genotype data includes ∼150,000 volunteers between 40-69 years old from the UK, genotyped at approximately 820,967 single nucleotide polymorphisms (SNPs). 50,000 samples were genotyped using the UKBiLEVE array and the remaining samples were genotyped using the UK Biobank Axiom array (Affymetrix)^84,85^. The UK Biobank Axiom is an update of UKBiLEVE and the two arrays share 95% of their content. In total, after sample and SNP quality control (QC), which was carried out centrally, 152,763 individuals and 806,466 directly typed SNPs remained. Phasing, imputation and derivation of principal components were also carried out centrally. Following imputation the number of variants reached 73,355,667 in 152,249 individuals. We performed additional quality control (QC) checks. We excluded samples with call rate ≤97%. We checked samples for gender discrepancies, excess heterozygosity, relatedness, ethnicity and we removed possibly contaminated and withdrawn samples. Following QC, the number of individuals was 138,997. We excluded 528 SNPs that had been centrally flagged as subject to exclusion due to failure in one or more additional quality metrics.

To define OA cases, we used the self-reported status questionnaire and the Hospital Episode Statistics data (Supplementary Table 3). We conducted five OA discovery GWAS and one sensitivity analysis, and the case strata were: self-reported OA at any site n=12,658; sensitivity analysis (a random subset of the self-reported cohort equal to the sample size of the hospital diagnosed cohort) n=10,083; hospital-diagnosed OA at any site based on ICD10 and/or ICD9 hospital records codes n=10,083; hospital-diagnosed hip OA n=2,396; hospital-diagnosed knee OA n= 4,462; and hospital-diagnosed hip and/or knee OA n=6,586. We applied exclusion criteria to minimise misclassification in the control datasets to the extent possible (using approximately 4x the number of cases for each definition) (Supplementary Table 2, Supplementary Fig. 1). We restricted the number of controls used and did not utilise the full set of available genotyped control samples from UK Biobank in order to guard against association test statistics behaving anti-conservatively in the presence of stark case: control imbalance for alleles with MAC <400^26^ (analogous to MAF ∼0.02 in the self-reported and hospital diagnosed OA datasets). For the control set, we excluded all participants diagnosed with any musculoskeletal disorder, or with relevant symptoms or signs, such as pain and arthritis, and selected older participants to ensure we decrease the number of controls that might be diagnosed with OA in the future, while keeping the number of males and females balanced (Supplementary Table 1).

At the SNP level, we further filtered for Hardy Weinberg equilibrium (HWE) *P*≤10^−6^, MAF≤0.001 and info score<0.4 (Supplementary Fig. 1). We tested for association using the frequentist LRT test and method ml in SNPTEST v2.5.2^86^ with adjustment for the first 10 principal components in order to control for population structure. Power calculations were carried out using Quanto v1.2.4^87^.

### Replication

Two hundred independent and novel variants with *P*<1.0×10^−5^ in the discovery analyses were taken forward for *in silico* replication in an independent cohort from Iceland (deCODE) using fixed effects inverse-variance weighted meta-analysis in METAL^88^. One hundred and seventy three variants were present in the replication cohort. The deCODE dataset comprised four OA phenotypes: any OA site (18,069 cases and 246,293 controls), hip OA (5,714 cases and 199,421 controls), knee OA (4,672 cases and 172,791 controls), and hip and/or knee OA (9,429 cases and 199,421 controls). We performed meta-analyses (across OA definitions) using summary statistics from the UK Biobank OA analyses and deCODE. We use *P*≤5x10^−8^ as the threshold to report genome wide significance.

*Replication cohort*: The information on hip, knee and vertebral osteoarthritis was obtained from Landspitali University Hospital electronic health records, Akureyri Hospital electronic health records and from a national Icelandic hip or knee arthroplasty registry^89^. Secondary osteoarthritis (e.g. Perthes disease, hip dysplasia), post-trauma osteoarthritis (e.g. ACL rupture) and those also diagnosed with rheumatoid arthritis were excluded from these lists. Only those diagnosed with osteoarthritis after the age of 40 were included. Hand osteoarthritis patients were drawn from a database of 9,000 hand osteoarthritis patients that was initiated in 1972^24^. The study was approved by the Data Protection Authority of Iceland and the National Bioethics Committee of Iceland. Informed consent was obtained from all participants.

### Association with OA-related endophenotypes

The 9 replicating genetic loci were examined for association in radiographic OA endophenotypes. This was done for 3 phenotypes: minimal Joint space width (mJSW), and two measures of hip shape deformities known as strong predictors for OA: acetabular dysplasia (measured with Center Edge-angle), and cam deformity (as measured with alpha angle)^90^. For mJSW association statistics for the variants were looked-up in a previously published GWAS, which joint analyzed data from the Rotterdam Study I (RS-I), Rotterdam Study II (RS-II), TwinsUK, SOF and MrOS using standardized age, gender and population stratification (four principal components) adjusted residuals from linear regression^50^. For the two hip shape phenotypes, CE-angle and alpha angle were measured as previously published^90^. CE-angle was analyzed as a continuous phenotype. We conducted GWAS on a total of 6880 individuals from the Rotterdam Study I (RS-I), Rotterdam Study II (RS-II), Rotterdam Study III (RS-III) and CHECK datasets using standardized age, gender adjusted residuals from linear regression. For cam-deformity individuals with an alpha-angle >60° were defined as a case (n=639), while all others were controls (4339). The GWAS was done on individuals from RS-I, RS-II and CHECK, using age and sex as covariates. The results of the discovery cohorts were quality checked using EASYQC^91^. The results of the separate cohorts were combined in a meta-analysis using inverse variance weighting with METAL^88^. Genomic control correction was applied to the standard errors and *P-values* before meta-analysis.

### Heritability estimation

To investigate the narrow sense heritability and the genetic correlation between the five osteoarthritis disease definitions, we ran the LDscore^92^ method that uses summary statistics at common-frequency variants genome-wide (independent of *P-value* thresholds) and LD estimates between variants while accounting for sample overlap. To estimate heritability on the liability scale, the sample prevalence was set to 20%. To calculate the population prevalence in the UK (65 million people), we consulted Arthritis Research UK figures: 8.75 million people have symptomatic osteoarthritis, while 2.46 and 4.11 million people have osteoarthritis of the hip and the knee, respectively. We assumed that 2.46+4.11 million people have osteoarthritis of the hip and/or the knee. To assess whether the osteoarthritis heritability estimates from self-reported (h1) and medical records (h2) were significantly different, we assumed h1-h2 ∼ N(h1-h2, variance(h1) + variance(h2) -2*rho*se(v1)*se(v2)), where rho is the correlation between h1 and h2, which was approximated by the Pearson’s product-moment correlation and the Spearman’s rank correlation of the negative log of the *P-values* and ORs when restricting analyses to common variants.

### Functional genomics

#### Patients and samples

We collected cartilage samples from 38 patients undergoing total joint replacement surgery: 12 knee OA patients (cohort 1; 2 women, 10 men, age 50-88 years); 17 knee OA patients (cohort 2; 12 women 5 men, age 54-82 years); 9 hip OA patients (cohort 3; 6 women, 3 men, age 44-84 years). We collected matched intact and degraded cartilage samples from each patient (see below). Cartilage was separated from bone and chondrocytes were extracted from each sample. From each isolated chondrocyte sample, we extracted RNA and protein. All patients provided full written informed consent prior to participation. All sample collection, RNA and protein extraction steps are described in detail in^93^.

#### Cohorts 1 and 2

This work was approved by Oxford NHS REC C (10/H0606/20), and samples were collected under Human Tissue Authority license 12182, Sheffield Musculoskeletal Biobank, University of Sheffield, UK. We excluded any patients with diagnosis other than osteoarthritis, with a history of glucocorticoid use (systemic or intra-articular) within the previous 6 months, or use of any other drugs associated with immune modulation; we further excluded patients with any history of fracture, significant knee surgery (apart from meniscectomy), knee infection, or any malignancy within the previous 5 years. Samples were scored using the OARSI cartilage classification system^94,95^: one sample with high OARSI grade signifying high-grade degeneration (“degraded sample”), and one sample with low OARSI grade signifying healthy tissue or low-grade degeneration (“intact sample”). Intact tissue here denotes OA early in the degeneration process, derived from the same background disease joint environment, enabling a comparison with highly degraded tissue to understand OA pathogenesis processes.

#### Cohort 3

Samples were collected under National Research Ethics approval reference 11/EE/0011, Cambridge Biomedical Research Centre Human Research Tissue Bank, Cambridge University Hospitals, UK. Osteoarthritis status was confirmed by examination of the excised femoral head. Samples were classified macroscopically and visually as high-grade (“degraded”) or low-grade (“intact”).

*Proteomics*: Proteomics analysis was performed on intact and degraded cartilage samples from 24 individuals (15 from cohort 2, 9 from cohort 3). LC-MS analysis was performed on the Dionex Ultimate 3000 UHPLC system coupled with the Orbitrap Fusion Tribrid Mass Spectrometer. To account for protein loading, abundance values were normalised by the sum of all protein abundances in a given sample, then log2-transformed and quantile normalised. We restricted the analysis to 3917 proteins that were quantified in all samples. We tested proteins for differential abundance using limma^96^ in R, based on a within-individual paired sample design. Significance was defined at 1% Benjamini-Hochberg False Discovery Rate (FDR) to correct for multiple testing. Of the 3732 proteins with unique mapping of gene name and Ensembl ID, we took forward 245 proteins with significantly different abundance between intact and degraded cartilage at 1% FDR.

#### RNA sequencing

We performed a gene expression analysis on samples from all 38 patients. Multiplexed libraries were sequenced on the Illumina HiSeq 2000 (75bp paired-end read length). This yielded bam files for cohort 1 and cram files for cohorts 2 and 3. The cram files were converted to bam files using samtools 1.3.1^97^ and then to fastq files using biobambam 0.0.191^98^, after exclusion of reads that failed QC. We obtained transcript-level quantification using salmon 0.8.2^99^ (with -gcBias and -seqBias flags to account for potential biases) and the GRCh38 cDNA assembly release 87 downloaded from Ensembl [<u>http://ftp.ensembl.org/pub/release-87/fasta/homo_sapiens/cdna/</u>]. We used tximport^100^ to convert transcript-level to gene-level count estimates, with estimates for 39037 genes based on Ensembl gene IDs. After quality control, we retained expression estimates for 15994 genes with counts per million of 1 or higher in at least 10 samples. Limma-voom^101^ was used to remove heteroscedascity from the estimated expression data. We tested genes for differential expression using limma^96^ in R (with lmFit and eBayes), based on a within-individual paired sample design. Significance was defined at 1% Benjamini-Hochberg False Discovery Rate (FDR) to correct for multiple testing. Of the 14408 genes with unique mapping of gene name and Ensembl ID, we took forward 1705 genes with significantly different abundance between intact and degraded cartilage at 1% FDR.

### Fine-mapping

We constructed regions for fine-mapping, by taking a window of at least 0.1 centimorgans either side of each index variant. The region was extended to the furthest variant with r^2^>0.1 with the index variant within a 1Mb window. LD calculations for extending the region were based on whole-genome sequenced EUR samples from the combined reference panel of UK10K^102^ and 1000 Genomes Projects^103,104^. For each region we implemented the Bayesian fine-mapping method CAVIARBF^105^, which uses association summary statistics and correlations among variants to calculate Bayes’ factors and posterior probabilities of each variant being causal. We assumed a single causal variant in each region and calculated 95% credible intervals, which contain the minimum set of variants that jointly have at least 95% probability of including the causal variant. We also applied the extended CAVIARBF method that uses functional annotation scores to upweigh variants according to their predicted functional scores. To this end, we downloaded pre-calculated CADD^106^ and Eigen^107^ scores from their equivalent websites. We observed better separation of severe-consequence genic variants with the CADD score and better separation of regulatory variants with the Eigen score, and therefore created a combined score, where splice acceptor, splice donor, stop lost, stop gained, missense and splice region variants were assigned their CADD-Phred score and the rest their Eigen-Phred score.

### Functional enrichment analysis

We used genome-wide summary statistics to test for enrichment of functional annotations. We used GARFIELD^108^ with customized functional annotations, making use of the functional genomics data we generated in primary articular chondrocytes using RNA sequencing and quantitative proteomics. We defined differentially transcribed genes and differentially expressed genes when comparing intact to degraded cartilage (1% FDR). We extended each differentially regulated gene by 5kb each side. Using GARFIELD’s approach, we calculated the effective number of independent annotations to be 1.995, which led to an adjusted p-value significance level of 0.025. We tested for enrichment using variants with *P*<1.0x10^−5^ and no analysis surpassed the corrected significance threshold.

### Pathway analysis

Gene-based and gene-set analyses were performed using MAGMA v1.06^109^ and consisted of three steps. First, each gene was assigned the variants located between its start and stop sites based on NCBI 37.3 gene definitions. Second, gene-based analysis was carried out on variant *P-values* derived from the GWAS results of each phenotype. Genotype data of 10,000 controls (subset of the 50,898 controls used in the self-reported OA analysis) were used to estimate linkage disequilibrium. *P-values* generated by MAGMA are not corrected for multiple testing. The genome-wide significance threshold for gene-based associations was calculated using the Bonferroni method (α = 0.05/number of genes being analysed). Genes containing 10 or fewer annotated variants were not included in the analyses. Third, a competitive gene-set analysis was carried out, implemented as a linear regression model on a gene data matrix which is created internally from the gene-based results. We used Kyoto Encyclopedia of Genes and Genomes^110^ (KEGG) and Reactome^111^ ^112^ Reactome: a knowledgebase of biological pathways gene annotations downloaded from MSigDB^113^ (version 5.2) on 23 January 2017. We also downloaded Gene Ontology^114^ (GO) biological process and molecular function gene annotations from Ensembl^115^ (version 87). We used annotations with the following evidence codes: a)Inferred from Mutant Phenotype (IMP) b) Inferred from Physical Interaction (IPI) c) Inferred from Direct Assay (IDA) d) Inferred from Expression Pattern (IEP) and e) Traceable Author Statement (TAS). KEGG/Reactome and GO annotations were analysed separately and only pathways that contained between 20 and 200 genes were included (595 for KEGG/Reactome, 619 for GO). Both self-contained (indicates that a gene-set does contain some associated genes) and competitive (assesses whether a pathway is more associated with a trait than other pathways) tests were applied, and we corrected for the potentially confounding effects of sample size, gene size, gene density and the inverse of the mean minor allele count (MAC) in the gene, as well as the log of these variables. MAGMA provides a built-in family-wise error rate (FWER) correction method. 10,000 permutations were used in each analysis and the significance threshold was set at α=0.05. We used DAPPLE^116^ for visualization of the pathways and protein-protein interaction (PPIs) relationships among the genes in each gene-set by integrating data from the InWeb database^117,118^ (Supplementary Fig. 6).

### LD regression

We used LDHub^119^ [accessed 23-27 January] to estimate the genome-wide genetic correlation between each of the OA definitions and 219 other human traits and diseases. In each analysis, we extracted variants with rsIDs (range 11,999,363-15,561,966) and uploaded the corresponding association summary statistics to LDHub, yielding 896,076-1,172,130 variants overlapping with LDHub. We corrected for multiple testing by defining significance at 5% Benjamini-Hochberg False Discovery Rate (FDR) for each of the five OA analyses.

### Mendelian randomization analysis

We used Mendelian randomization (MR) to assess the potential causal role of the phenotypes identified in the LD score regression analysis on osteoarthritis. We also included birth weight and height (Supplementary Table 21). MR uses genetic variants robustly associated with a given disease risk factor or exposure phenotype of interest as instrumental variables, in order to estimate the causal effect of such exposure phenotype on disease risk. MR yields valid causal effect estimates if the genetic instruments are: i) associated with the exposure phenotype; ii) not associated with exposure-outcome confounders; iii) not associated with the outcome except through the exposure. If the genetic instrument influences the disease outcome through pathways not fully mediated by the exposure phenotype (i.e. through horizontal pleiotropic effects), then the causal effect estimate will be biased. We applied a range of sensitivity analyses that allow relaxation of the assumptions of no horizontal pleiotropy in different ways (see below). In all analyses, the primary outcome variable was self-reported osteoarthritis. We used data from hospital records (which were available for a much smaller number of individuals) as sensitivity analyses and to identify potential site-specific effects.

#### Data sources

Genetic instruments were identified from publicly-available summary GWAS results through the TwoSampleMR R package, which allows extracting the data available in the MR-Base database^120^. Only results that combined both sexes were extracted. Preference was given to studies restricted to European populations to minimise the risk of bias due to population stratification; however, for a few traits those were either not available or corresponded to much smaller studies (Supplementary Table 21). However, this is unlikely to substantially bias the results because all studies employed correction methods, and even multi-ethnic studies are mostly composed of European populations. The exception was for number of children ever born and age of the individual when his/her first child was born: given that the GWAS of reproductive traits by Barban and colleagues^121^ was not available in MR-Base, we extracted summary association results for the variants that achieved genome-wide significance directly from the paper, and used coefficients from each sex in sex-specific analyses. The search was performed on June 19, 2017. For each trait, all genetic instruments achieved the conventional levels of genome-wide significance (i.e., *P*<5.0x10^−8^) and were mutually independent (i.e. *r^2^*<0.001 between all pairs of instruments).

#### Two-sample MR

For the exposure phenotypes with at least one genetic instrument available, we used two-sample MR analysis to evaluate their causal effect on osteoarthritis risk. The exceptions were smoking and reproductive traits, which were performed using one-sample MR only due to the need of performing the analysis within specific subgroups. All summary association results used for two-sample MR are shown in Supplementary Tables 22 and 23 provides an overall description of each set of genetic instruments. The two-sample design allows combining instrument-exposure summary association results from large GWAS consortia with the respective instrument-osteoarthritis summary association results from UK Biobank, thus minimising the possibility of weak instrument bias. Moreover, even if there is weak instrument bias, it attenuates (rather than exacerbates) the causal effect estimate^122^. Another strength of the two-sample design is that it allows implementing methods that make different assumptions regarding horizontal pleiotropy, thus partially relaxing the assumptions required for valid causal inference^123^. We applied the following methods:

- Ratio method: for exposure phenotypes with only one genetic instrument available, MR was performed using the ratio method, which consists of dividing the instrument-outcome by the instrument-exposure regression coefficient. The standard error of the ratio estimate can be calculated by dividing the instrument-outcome standard error by the instrument-exposure regression coefficient^124,125^. Confidence intervals and *P-value* were calculated using the Normal approximation.
- Inverse-variance weighting (IVW): this method consists of a weighted (by the inverse-variance of the instrument-outcome associations) linear regression of the instrument-outcome coefficients on the instrument-exposure coefficients, constraining the intercept to be zero (which follows from the no horizontal pleiotropy assumption^126^). We used a multiplicative random effects version of the method, which incorporates between-instrument heterogeneity in the confidence intervals^127^.
- MR-Egger regression: this is implemented as the IVW method, except that the intercept is not constrained. This method yields consistent causal effect estimates even if all instruments are invalid, provided that horizontal pleiotropic effects are uncorrelated with instrument strength (i.e. the Instrument Strength Independent of Direct Effects – InSIDE – assumption holds^128^).
- Weighted median: this method uses the median of the empirical inverse-variance weighted distribution function of all individual-instrument ratio estimates as the causal effect estimate. The weighted median estimate is a consistent causal effect estimate even if the InSIDE assumption is violated, provided that up to (but not including) 50% of the weights in the analysis come from invalid instruments^129^.
- Mode-based estimate (MBE): the MBE uses the mode of the empirical inverse-variance weighted smoothed density function of all individual-instrument ratio estimates as the causal effect estimate. The MBE relies on the ZEro Modal Pleiotropy Assumption (ZEMPA), which postulates that the largest subgroup (or the subgroup that carries the largest amount of weight in the analysis) of instruments that estimate the same causal effect estimate is composed of valid instruments. This allows consistent causal effect estimation even if the majority of instruments are invalid. The stringency of the method can be regulated by the *φ* parameter^130^. We tested two values of *φ*: *φ*=1 (ie, the default) and *φ*=0.5 (half of the default, or twice as stringent).

For exposure phenotypes with more than 1 but less than 10 genetic instruments, only the IVW method was applied. This was because the remaining methods are typically less powered and require a relatively large number of genetic instruments to provide reliable results. It possible to quantify the degree of weak instrument bias (which corresponds to regression dilution bias in two-sample MR) for the IVW and MR-Egger methods using the 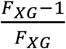 and 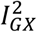 statistics, respectively. Both range from 0% to 100%, and 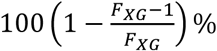 and 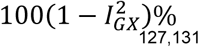 can be interpreted as the amount of dilution in the corresponding causal effect estimates^127,131^. Given that only genome-wide significant variants were selected as instruments, the 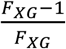 statistic will necessarily be high (approximately 95%, at least). However, the 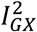 statistic depends both on instrument strength and heterogeneity between instrument-exposure associations, which implies that regression dilution bias in MR-Egger can be substantial even if instruments are individually strong. Indeed, for some traits the 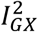 statistic was very low (Supplementary Table 23). Therefore, all MR-Egger regression analyses were corrected for regression dilution bias using a Simulation Extrapolation (SIMEX) approach^131^.

#### Horizontal pleiotropy tests

We additionally assessed the robustness of our findings to potential violations of the assumption of no horizontal pleiotropy by applying two tests of horizontal pleiotropy. One of them was the MR-Egger intercept, which can be interpreted as the average instrument-outcome coefficient when the instrument-exposure coefficient is zero. If there is no horizontal pleiotropy, the intercept should be zero. Therefore, the intercept provides an indication of overall unbalanced horizontal pleiotropy^128^. The second test was Cochran’s Q test of heterogeneity, which relies on the assumption that all valid genetic instruments estimate the same causal effect^132^.

#### Power calculations

We performed power calculations to estimate the power of our two-sample MR analysis to detect odds ratios of 1.2, 1.5 and 2.0. The analysis were performed through simulations (10,000 iterations), with power defined as the proportion of times that 95% confidence intervals of the IVW method excluded 1. Continuous exposure phenotypes were analysed in standard deviation units (e.g. odds ratio of OA of 1.2 per standard deviation increment in the exposure). Binary exposure phenotypes, were analysed in 0.3438196×ln(odds ratio) units (e.g. odds ratio of OA of 1.2 per 0.3438196 increments in the ln(odds ratio) of exposure), which corresponds to an average absolute increment of 0.5 in the odds ratio of exposure in an odds ratio range of 1.0 to 2.0. In each simulation, each instrument-exposure association was sampled from a Normal distribution 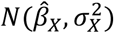, where 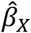 denotes the observed instrument-exposure regression coefficient, and *σ*_*X*_ is the observed instrument-exposure standard error. The corresponding instrument-outcome association was sampled from a Normal distribution 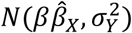, where *β* denotes the causal effect of the exposure phenotype on osteoarthritis (e.g. ln(1.2)), and *σ*_*Y*_ is the observed instrument-exposure standard error. This simulation model assumes that all genetic instruments are valid and estimate the same causal effect.

#### One-sample MR

UK Biobank data were used to perform one-sample MR using the same genetic instruments than in the two-sample MR. This analysis comprised an additional sensitivity analysis, with results that were concordant results between the one-sample and two-sample approaches being considered more robust. Another motivation was to perform analyses that require stratification: in the case of smoking, the genetic instrument (i.e. the *CHRNA3* variant rs12914385^133^) is associated with heaviness of smoking among smokers, so no association between the genetic instrument and osteoarthritis is expected among never smokers (unless the instrument affects osteoarthritis through smoking-independent pathways); in the case of reproductive traits, the effects of many genetic instruments were sex-dependent^121^, so we performed the analyses within sexes. A third motivation for the one-sample MR was to obtain multiple testing corrected *P-values* through permutation. This analysis used standardised allele scores weighted by the instrument-exposure summary associations as instrumental variables. First, the association between each weighted allele score and OA was tested. This comprises a test for presence and direction of causality. For the cases that presented multiple testing corrected statistical evidence of association, the causal effect estimate per unit change in the exposure phenotype was estimated using a two-stage approach: in the first stage, genetically-predicted values of the exposure phenotype were generated by regressing the phenotype on its weighted allele score using linear regression; in the second stage, logistic regression was used to estimate the effect of the genetically-predicted values of the exposure phenotype on OA and, hence, the causal effect of the exposure on OA. To account for uncertainty in the prediction in the first-stage, standard errors of the causal effect estimate were bootstraped.

